# Metabolite Profiling and Cytotoxic Investigation of *Senecio biafrae* Oliv. & Hiern (Compositae) on Nauplii Brine Shrimps and Cancer Cell Lines

**DOI:** 10.1101/2023.05.21.541662

**Authors:** Funke Toyin Akinola, Olumayowa Vincent Oriyomi, Seun Bayonle Ogundele, Mukaila Babatunde Adekola, Morounfolu Joseph Agbedahunsi, Olusegun Olubunmi Babalola

**Author notes:** Corresponding Author Akinola, Funke Toyin, Biochemistry and Molecular Biology, Faculty of Science, Obafemi Awolowo University, Ile-Ife, Nigeria.

## Abstract

The study investigated the cytoactivities of methanolic leaf extract (MLE) and fractions of *Senecio biafrae*, profiled, and elucidated their metabolites. Hydromethanolic extract of *S. biafrae* obtained by maceration was partitioned and subjected to cytotoxicity using brine shrimp lethality assay (BSLA), human cervical adenocarcinoma (HeLa) and prostate cancer cell (PC3) lines. The bioactive principles of most active fractions were profiled using GC-MS. The dichloromethane fraction (DMF) was purified using column and high-performance liquid chromatography to afford compounds 1 and 2 identified by 1H and 13C, DEPT-90, DEPT-135, COSY, NOESY and EIMS spectra. The MLE and its fractions exhibited good toxicity on nauplii brine shrimp. The HF exhibited a pronounced cytoactivity on PC-3 cell line with 92.8% followed by DMF. The GC-MS profiling of HF and DMF identified 17 and 6 constituents respectively comprising saturated and unsaturated fatty acids, diterpenoidal alcohol, phytol and aromatic compounds. Hexadecanoic acid, cis, cis, cis-9, 12, 15-octadecatrienoic acid, and phytol accounted for the highest percentage of the constituents in both fractions. Structure elucidation confirmed compounds 1 and 2 as stigmasterol and ergosterol with cytoactivity which compared favourably with doxorubicin. The study revealed that *S*.*biafrae* contains cytotoconstituents evidenced by proliferation inhibition of HeLa and PC-3 cell lines. The cyto-activities and bioconstituents indicate that *S*.*biafrae* has the potential to be considered as a candidate for drug development in the management of cancer-related conditions. Further investigation into the specific bioactive compounds and their mechanisms of action could provide valuable insights into the therapeutic potential of *S*.*biafrae* in cancer treatment.

## INTRODUCTION

From time immemorial, Man have depended on nature for basic needs of life like food, shelter, clothing, etc. with majority of these natural products coming from plants, animals and minerals needed for treating human diseases ^[1, 2]^. Plants have formed the basis of sophisticated traditional medicine systems that have been in existence for thousands of years and continue to provide mankind with new remedies ^[3]^. This system continues today because of its biomedical benefits and cultural beliefs in many parts of the world which made it a great contribution towards maintaining human health ^[4]^. The World Health Organization estimated that 80% of the population in the developing countries continues to use traditional medicine to treat infectious diseases ^[5]^.

Medicinal plants are known to contain inherent active ingredients used to manage disease or relieve pain ^[6]^. The use of traditional medicines and medicinal plants as therapeutic agents for the maintenance of good health in most developing countries has been widely observed ^[7]^. According to World Health Organization, medicinal plants are the best sources for obtaining variety of drugs. Therefore, plants should be investigated to better understand their properties, safety and efficacy ^[8]^. The efficacy of medicinal plants in the management of diseases is indubitable as it occupies a unique place in human life and provides information about the use of plants or plant parts as medicine ^[9]^. *Senecio biafrae* Oliv. & Hiern (Compositae), commonly known as life root plant, is one of the most important medicinal plants with edible leaves and stems, widely consumed as vegetables among the Yoruba tribe of South-western, Nigeria ^[10]^. There is a resurgent of interest in the pharmacological efficacy of medicinal plants in the management of various human carcinomas and other pathologies, especially in the face of increasing failure and toxicity associated with the use of some anticancer agents ^[11]^. Numerous bioactive compounds or their derivatised complexes from medicinal plants have been reported for chemopreventive and anticancer activity ^[12]^. Phytochemical investigations of *Senecio biafrae* revealed the presence of phytometabolites belonging to different classes of chemical diversities such as terpenoids, flavonoids, alkaloids, anthocyanins, coumarins, glycosides and dihydroisocoumarins which have been reported to possess antimicrobial properties ^[13]^. The leaf of *S. biafrae* is ethnomedicinally used for the treatment of wounds, sore eyes, infertility, rheumatism, pulmonary defects, heart problems, cough and diabetes and promotion of milk secretion in lactating mothers ^[14-15]^. *S. biafrae* has been reported to be a good source of protein having high levels of essential amino acids and vitamins A, C and E, and has been recommended as a food supplement ^[16]^. In this study, we investigated the cytotoxic activity of *Senecio biafrae* leaf extract and its fractions on nauplii brine shrimp, human cervical adenocarcinoma (HeLa) and prostate cancer cell (PC-3) line**s** and carried out the chemical profiling, isolation, purification and structure elucidation of the active chemical principles on the two most active fractions.

## MATERIALS

### Chemicals and Reagents

Fetal bovine serum (FBS), trypan blue solution and antibiotics (streptomycin/penicillin) were purchased from Gibco (California, USA). RPMI-1640 medium, D-Glucose, HEPES, Dulbecco modified eagle medium (DMEM), Triton X100, F12-DMEM and 3-(4, 5-dimethylthiazol-2-yl)-2, 5-diphenyltetrazolium bromide (MTT) were obtained from Sigma-Aldrich (St. Louis, MO, USA). Dimethyl sulfuroxide (DMSO) was purchased from Fisher Scientific (Leicestershire, UK). *Escherichia coli* Lipopolysaccharide (LPS) was purchased from DIFCO Laboratories (Michigan, USA). PC-3 (human epithelial, prostate cancer) and HeLa (human epithelial adenocarcinoma) cells were obtained from European Collection of Cell Cultures, Salisbury ECACC (UK). HPLC-graded methanol, chloroform, hexane and ethyl acetate were purchased from Merck (Darmstadt, Germany). Tween 20, ethanol, pre-coated silica gel (F_254_) plates, silica gel (SiO_2_; 230-400 mesh), butanol, dichloromethane and chloroform were purchased from Merck, (Damstadt, Germany). De-ionized water was obtained from a Milli-Q UV-Plus water purification system (Millipore Corp., Billerica, MA, USA).

### Collection and Identification of Sample

Fresh leaves of *Senecio biafrae* were collected in January, 2019 from Akamo Village Iperindo, Osun State, Nigeria (Latitude 07°29’40”; Longitude E04°49’26”). The plant was identified and authenticated at the IFE Herbarium, Department of Botany, Obafemi Awolowo University, Ile – Ife, Nigeria. Voucher specimen (voucher number (IFE-17936) was prepared and deposited in the Herbarium of the department of Botany, Obafemi Awolowo University, Nigeria. The nomenclature of the plant was further confirmed at www.worldfloraonline.org.

## METHODS

### Extraction and Partitioning of Plant

Powdered leaf (2 kg) was macerated (3X) with 80% (v/v) MeOH: H_2_O (10 L) at room temperature for 48 h with continuous agitation on a shaker. The combined extract was then filtered and evaporated *in vacuo* at 40°C to afford a dry crude MeOH extract (45 g) which was subsequently suspended in H_2_O and successively partitioned to afford the respective fractions: *n*-hexane, dichloromethane (DCM), ethyl acetate (EtOAc), butanol and aqueous residue which were also concentrated to dryness *in vacuo* at 40°C. The crude extract and respective fractions were then tested for their cytotoxic activities. The n-hexane and dichloromethane fractions exhibited the most cytotoxic activity and were selected for further analysis.

### GC-MS Metabolite Profiling of Most Active Fractions

The metabolite profiling of fractions that exhibited the most cytoactivities was carried out in order to identify the chemical constituents responsible for their cytotoxic activity. They were analyzed on a fused silica AT-5MS capillary column (30m x 0.25mm internal diameter, 0.25 µm film thickness) equipped on a gas chromatograph Agilent 5975C combined with inert XL EICI MSD fitted with a triple-axis detector source (Agilent Technologies, Santa Clara, CA, USA). The split ratio of injection was set at 1:30 with injection pot temperature of 270°C, while the column temperature was set at 60°C upon injection. The temperature was increased at rate of 3°C/minute to 270°C and held for 30 minutes. Purified helium was the carrier gas at flow rate of 1mL/minute. The mass spectrometer was operated in EI mode at 70eV, ion source temperature of 230°C, quadrupole temperature 150°C, mass range (m/e) of 29-550 with scan rate of 6.35 scan/s and electronmultiplier voltage 1341V. The MS transfer line temperature was set at 280°C. Identification of components was performed based on their retention time and by mass spectra matching with National Institute of Standard and Technology (NIST) MS library 2009, while concentration of identified components was calculated using area normalization over flame ionization detector response.

### Chromatographic Fractionation of DCM Fraction

The DCM fraction (10g) was chromatographed and fractionated over silica gel on open column and eluted with *n*-hexane with increasing 10% gradient of EtOAc up to 100% EtOAc followed by increasing gradient elution with MeOH up to 100% MeOH to afford 13 fractions A-M. Sub-fraction G was further fractionated over sephadex LH 20 column and isocratically eluted with DCM:MeOH (9:1) to afford sub-fractions 1-30 of 10 mL each, which were analyzed by TLC. Sub-fractions 6-15 were combined and further chromatographed on silica gel column to afford Compound 1 as a white powder, which precipitated out and was recrystallized with CHCl_3_. Sub-fraction H was also chromatographed on open column chromatography and eluted with dichloromethane up to 100% followed by gradient with EtOAc up to 100% followed by gradient with MeOH to give sub-fractions H1-H10. Following TLC analysis, sub-fractions H6-H10 were combined and was further chromatographed on open column with silica gel 230-400 mesh as stationary phase and eluted with dichloromethane with increasing EtOAc up to 100% and MeOH solvent systems to afford an off-white powder, which was dissolved in CH_2_Cl_2_ and purified on sephadex column to afford Compound 2 which was recrystallized from CHCl_3_ to give a white solid.

### Brine Shrimp Lethality Bioassay

The extract and fractions of *S. biafrae* leaf were evaluated in a test for lethality to brine shrimp larvae according to Meyer *et al*. (1982). Toxicities of compounds were tested at 10, 100, and 1000 µg mL in 10 mL sea-water solutions with 1% DMSO (v/v). Ten nauplii were used in each test and survivors counted after 24 h. Three replications were used for each concentration. The blank control is conducted with Distilled water. The median lethal concentration of the fractions for 50% mortality after 24 h of exposure was calculated using Probit Analysis (Finney, 1971) using the mathematical expression, % mortality = (no. of dead nauplii/initial no. of live nauplli) × 100.

### *In vitro* Cell Culture

Human cervical adenocarcinoma and prostate cancer cell line**s** were used to evaluate the cytotoxic effect of extract and fractions while Compound 1 was investigated on HeLa cell line. The cells were grown in Dulbecco modified eagle medium (DMEM) (Sigma-Aldrich (St. Louis, MO, USA) supplemented with 10% fetal bovine serum and 1% penicillin/ streptomycin (Gibco, USA). Cells were maintained in a humidified atmosphere containing 5% CO_2_ at 37°C.

### *In vitro* Cytotoxicity Assay

The cytotoxic effect of extract, fractions and isolated compounds on PC3 and HeLa cells was evaluated by colorimetry assay using 3-(4, 5-dimethylthiazole-2-yl)-2, 5-diphenyl-tetrazolium bromide (MTT) ^[17].^ Extract, fractions and isolated compounds were individually dissolved in 1% DMSO and diluted with cell culture medium to give final concentration of 30 µg/mL. A 100 µL of HeLa and PC3 cells suspensions (6×10^4^ cells/mL) was seeded in a 96-well plate and incubated overnight at 37^°^C in 5% CO_2_. The medium was carefully aspirated and the cells were challenged with the prepared solutions of extract, fractions and isolated compounds and the plate was further incubated for further 48 hours. An aliquot of 50 µL MTT at concentration of 0.5 mg/mL was then dispensed into each well and incubated for 4 hours, followed by the addition of 100 µL of DMSO. The extent of MTT reduction to formazan within cells was calculated by measuring the absorbance at 570 nm using a micro-plate reader (Spectra Max Plus, Molecular Devices, CA, USA). The cytotoxicity was recorded as percentage inhibition of the sample and standard on the cell lines. Doxorubicin (standard drug) was used as a positive control, while DMSO served as negative control. The percent inhibition was calculated by using the following formula:

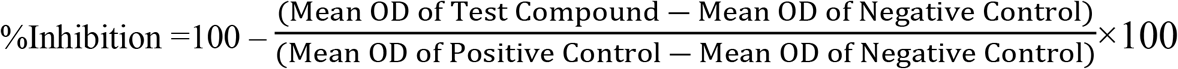

### Statistical Analysis

Data obtained were expressed as mean ± S.E.M using Graph Pad Instat Graphical Statistical Package version 5 (GraphPad Software Inc, USA). Significance of the results were evaluated using one-way analysis of variance (ANOVA) and the means was compared using Turkey’s test. Values of p < 0.05 was considered as statistically significant

## 3. RESULTS

### Brine Shrimp Lethality Assay of *S. biafrae* Plant

The result of brine lethality assay of *S. biafrae* extract and fractions is represented in Table 1.

**Table 1:** Percentage Mortality of shrimp nauplii after treating with *Senecio biafrae* extract and fractions.

| Plant Extract | Concentration<br>(µg mL) | Number of Surviving Nauplii<br>(after 24 h) |  |  | Total Number of<br>Nauplii Survivor | %<br>Mortality | LC50<br>(µg/mL) |
| --- | --- | --- | --- | --- | --- | --- | --- |
|  |  | R1 | R2 | R3 |  |  |  |
| Control<br>(Distilled water) | 10 | 10 | 10 | 10 | 30 | 0% | 0 |
|  | 100 | 10 | 10 | 10 | 30 | 0% |  |
|  | 1000 | 10 | 10 | 10 | 30 | 0% |  |
| Standard<br>(Etoposide) | 10 | 10 | 9 | 9 | 28 | 6.67% | 465.64 |
|  | 100 | 7 | 6 | 6 | 19 | 36.67% |  |
|  | 1000 | 1 | 1 | 0 | 2 | 93.33% |  |
| *S. bialfrae*<br>(Methanolic Extract) | 10 | 10 | 10 | 9 | 29 | 3.33% | 571.32 |
|  | 100 | 10 | 10 | 9 | 29 | 3.33% |  |
|  | 1000 | 9 | 9 | 9 | 27 | 10% |  |
| *S. bialfrae*<br>(Hexane Fraction) | 10 | 10 | 10 | 10 | 30 | 0% | 610.74 |
|  | 100 | 10 | 10 | 9 | 29 | 3.33% |  |
|  | 1000 | 3 | 3 | 3 | 9 | 70% |  |
| *S. bialfrae*<br>(DCM Fraction) | 10 | 10 | 10 | 10 | 30 | 0% | 610.75 |
|  | 100 | 10 | 9 | 9 | 28 | 6.66% |  |
|  | 1000 | 0 | 0 | 0 | 0 | 100% |  |
| *S. bialfrae*<br>(EA Fraction) | 10 | 10 | 10 | 10 | 30 | 0% | 572.73 |
|  | 100 | 10 | 9 | 9 | 28 | 6.66% |  |
|  | 1000 | 9 | 9 | 9 | 27 | 10% |  |
| *S. bialfrae*<br>(Butanol Fraction) | 10 | 9 | 9 | 9 | 27 | 10% | 483.98 |
|  | 100 | 8 | 8 | 8 | 24 | 20% |  |
|  | 1000 | 8 | 8 | 8 | 24 | 20% |  |
| *S. bialfrae*<br>(Aqueous Fraction) | 10 | 10 | 10 | 10 | 30 | 0% | 576.95 |
|  | 100 | 10 | 9 | 9 | 28 | 6.66% |  |
|  | 1000 | 8 | 8 | 8 | 24 | 20% |  |

### Cytotoxicity of *S. biafrae*

The cytotoxic activity of extract, fractions and isolated compounds on PC-3 and HeLa cell lines are presented in Table 2. The hexane showed better activity than the crude extract, and other fractions tested on the PC-3 cell lines. The cytotoxic activity revealed that *n*-hexane fraction of the methanolic extract *S. biafrae* leaf exhibited the best cytotoxic activity with % inhibition of 92.8 on PC-3 cell lines and 20.2 on HeLa cell lines, and therefore is the most active fraction of the crude extract. Thus, in this study, the hexane fraction particularly demonstrated a better cytotoxic activity on PC-3 cell line followed by doxorubicin, dichloromethane, compounds 1 and 2. Compound 1 was found to exhibit 45.9 and 21.7 % inhibition on the proliferation of HeLa and PC-3 cell lines respectively compared to doxorubicin with 89.9 and 100 % inhibition at the same concentration of 30 µg/mL. Also compound 1 exhibited a better cytotoxic activity than compound 2.

**Table 2:** Cytotoxic Activity of Extract, Fractions and Compounds 1, 2 on PC-3 and HeLa cell.

| Fraction/Drug | PC3 | HeLa |
| --- | --- | --- |
|  | % Inhibition at 30µg/ml | % Inhibition at 30µg/ml |
| **Crude extract** | 0.73 | 6.9 |
| **Hexane** | 92.8 | 20.2 |
| **DCM** | 17.1 | 6.8 |
| **Ethyl acetate** | -17.7 | -8.4 |
| **Butanol** | -24.1 | 6.11 |
| **Aqueous** | -12.0 | -0.6 |
| **Compound 1** | 21.9 | 45.9 |
| **Compound 2** | 13.4 | 15.2 |
| **Doxorubicin** | 89.9 | 101.2 |
| **DMSO** | 0 | 0 |

### GC-MS Metabolite Profiling of Hexane and Dichloromethane Fractions

The GC-MS profiling of the most active fractions of methanolic extract identified seventeen (17) bioactive compounds in the hexane and six (6) in the dichloromethane fractions respectively. The total ion chromatogram is presented in Figures 1 and 2 while Tables 3 and 4 show the identified constituents, their names, molecular weight, molecular formula, retention time and abundance (peak area in percentage). The major compounds identified were n-Hexadecanoic acid (35.90%), 9, 12, 15-Octadecatrienoic acid, (Z, Z, Z) (21.85%), Phytol (9.40%), Hexadecanoic acid, methyl ester (7.38%), 9, 12-Octadecadienoic acid (Z,Z)-(6.08%) and 9,12,15-Octadecatrienoic acid, methyl ester, (Z,Z,Z)- (5.97%) while the minor compounds identified were 9,12-Octadecadienoic acid (Z,Z)-, methyl ester (2.79%), Octadecanoic acid (1.68%), 2-Pentadecanone, 6,10,14-trimethyl (1.52%), Octadecanoic acid, methyl ester (0.97%), 3,7,11,15-Tetramethyl-2-hexadecen-1-ol (0.69%), 1,2-Benzenedicarboxylic acid, diisooctyl ester (0.59%), 3,7,11,15-Tetramethyl-2-hexadecen-1-ol (0.46%), 2H-Pyran-2-one, tetrahydro-6-nonyl (0.32%), Eicosanoic acid, methyl ester (0.28%), 1,4-Benzenediol, 2,6-bis(1,1-dimethylethy (0.11%) and 3-Butoxy-1,1,1,5,5,5-hexamethyl-3-(trimethylsiloxy)trisiloxane (0.09%).

**Table 3:** Constituents Identified in Hexane Fraction of *S. biafrae* Leaf Extract. MW= Molecular weight, RT= Retention time

**Figure 2: GC-MS Chromatogram of Dichloromethane Fraction**
| S/N | Compounds | Formula | MW | RT(min) | Area |
| --- | --- | --- | --- | --- | --- |

|  |  |  |  |  | (%) |
| --- | --- | --- | --- | --- | --- |
| 1 | 2-Pentadecanone, 6,10,14-trimethyl | C<sub>18</sub>H<sub>36</sub>O | 268 | 35.01 | 1.52 |
| 2 | Hexadecanoic acid, methyl ester | C<sub>17</sub>H<sub>34</sub>O<sub>2</sub> | 270 | 41.00 | 7.38 |
| 3 | n-Hexadecanoic acid | C<sub>16</sub>H<sub>32</sub>O<sub>2</sub> | 256 | 43.90 | 35.90 |
| 4 | 9,12-Octadecadienoic acid (Z,Z)-, methyl ester | C<sub>19</sub>H<sub>34</sub>O<sub>2</sub> | 294 | 48.70 | 2.79 |
| 5 | 9,12,15-Octadecatrienoic acid, methyl ester, (Z,Z,Z)- | C<sub>19</sub>H<sub>32</sub>O<sub>2</sub> | 292 | 48.87 | 5.97 |
| 6 | Phytol | C<sub>20</sub>H<sub>40</sub>O | 296 | 49.27 | 9.40 |
| 7 | Octadecanoic acid, methyl ester | C<sub>19</sub>H<sub>38</sub>O<sub>2</sub> | 298 | 49.51 | 0.97 |
| 8 | 9,12-Octadecadienoic acid (Z,Z)- | C<sub>18</sub>H<sub>32</sub>O<sub>2</sub> | 280 | 49.73 | 6.08 |
| 9 | 9,12,15-Octadecatrienoic acid, (Z,Z,Z)- | C<sub>18</sub>H<sub>30</sub>O<sub>2</sub> | 278 | 49.91 | 21.85 |
| 10 | Octadecanoic acid | C<sub>18</sub>H<sub>36</sub>O<sub>2</sub> | 284 | 50.32 | 1.68 |
| 11 | 3,7,11,15-Tetramethyl-2-hexadecen-1-ol | C<sub>20</sub>H<sub>40</sub>O | 296 | 51.52 | 0.69 |
| 12 | Eicosanoic acid, methyl ester | C<sub>21</sub>H<sub>42</sub>O<sub>2</sub> | 326 | 53.40 | 0.28 |
| 13 | 2H-Pyran-2-one, tetrahydro-6-nonyl- | C<sub>14</sub>H<sub>26</sub>O<sub>2</sub> | 226 | 53.90 | 0.32 |
| 14 | 3,7,11,15-Tetramethyl-2-hexadecen-1-ol | C<sub>20</sub>H<sub>40</sub>O | 296 | 54.39 | 0.46 |
| 15 | 3-Butoxy-1,1,1,5,5,5-hexamethyl-3-(trimethylsiloxy)trisiloxane | C<sub>13</sub>H<sub>36</sub>O<sub>4</sub>Si<sub>4</sub> | 368 | 56.26 | 0.09 |
| 16 | 1,2-Benzenedicarboxylic acid, diisooctyl ester | C<sub>24</sub>H<sub>38</sub>O<sub>4</sub> | 390 | 56.61 | 0.59 |
| 17 | 1,4-Benzenediol, 2,6-bis(1,1-dimethylethy | C<sub>14</sub>H<sub>22</sub>O<sub>2</sub> | 222 | 56.93 | 0.11 |

**Table 4:** Metabolite Constituents Identified in the Dichloromethane Fraction of *S. biafrae*.

| S/N | Compounds | Formula | MW | RT(min) | Area (%) |
| --- | --- | --- | --- | --- | --- |
| 1 | 3,7,11,15-Tetramethyl-2-hexadecen-1-ol | C<sub>20</sub>H<sub>40</sub>O | 296 | 34.63 | 4.70 |
| 2 | n-Hexadecanoic acid | C<sub>16</sub>H<sub>32</sub>O<sub>2</sub> | 256 | 43.63 | 17.24 |
| 3 | Phytol | C<sub>20</sub>H<sub>40</sub>O | 296 | 49.28 | 12.67 |
| 4 | 9,12-Octadecadienoic acid (Z,Z)- | C<sub>18</sub>H<sub>32</sub>O<sub>2</sub> | 280 | 49.67 | 9.52 |
| 5 | 9,12,15-Octadecatrienoic acid, methyl ester, (Z,Z,Z)- | C<sub>19</sub>H<sub>32</sub>O<sub>2</sub> | 292 | 49.87 | 52.35 |
| 6 | 9,12,15-Octadecatrienoic acid, (Z,Z,Z)- | C<sub>18</sub>H<sub>30</sub>O<sub>2</sub> | 278 | 50.29 | 3.51 |

**Figure 1:**
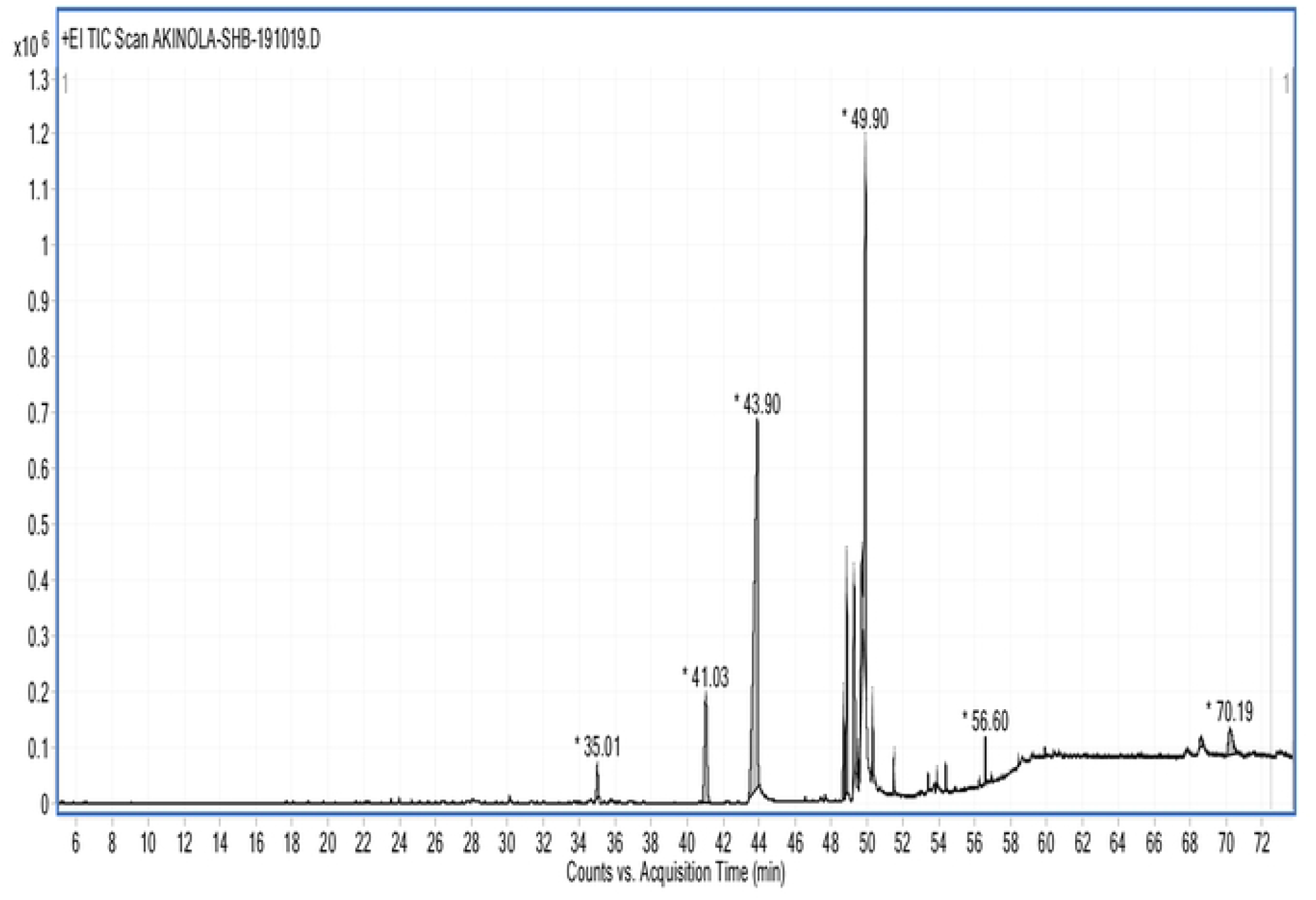
GC-MS Chromatogram of Hexane Fraction.

**Figure 2:**
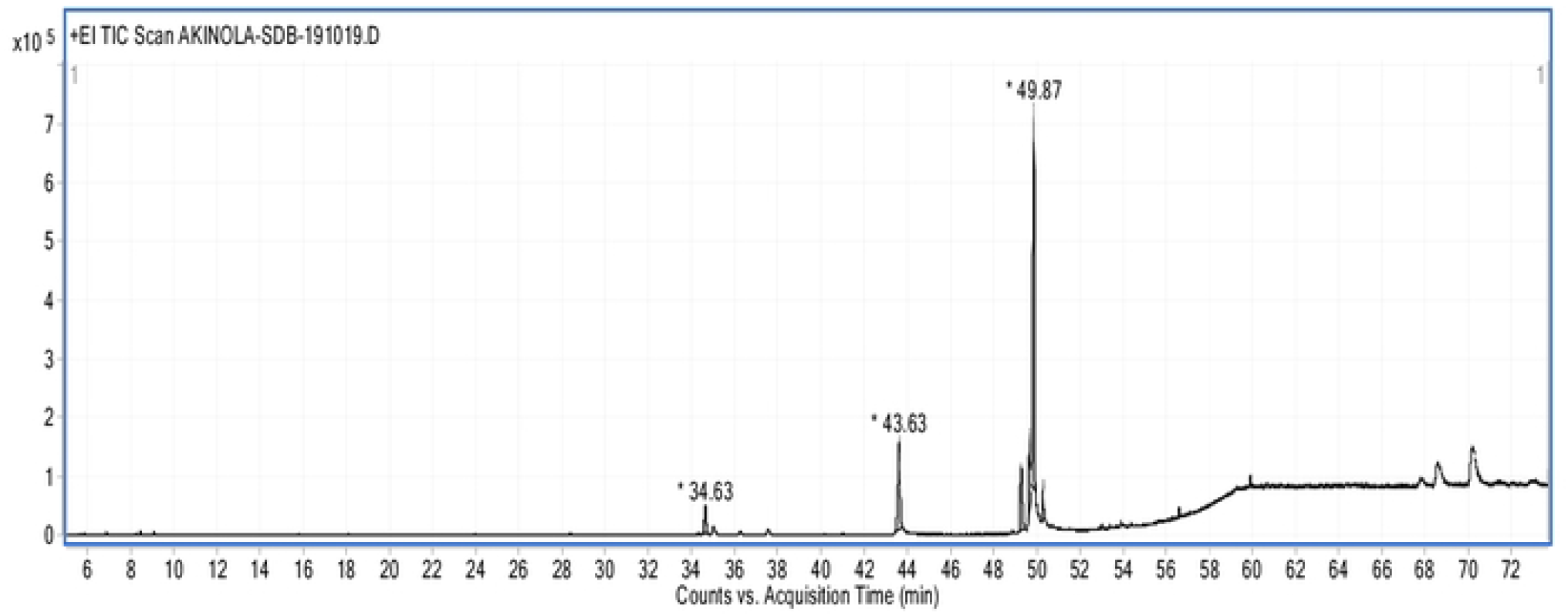
GC-MS Chromatogram of Dichloromethane Fraction.

### Chromatographic Fractionation of Dichloromethane (DCM) Fraction

Chromatographic fractionation to isolate and purify the chemical components in the DCM fraction afforded two known triterpenoids.

Compound 1 (Stigmasterol) Figure 3

**Figures 3 and 4:**
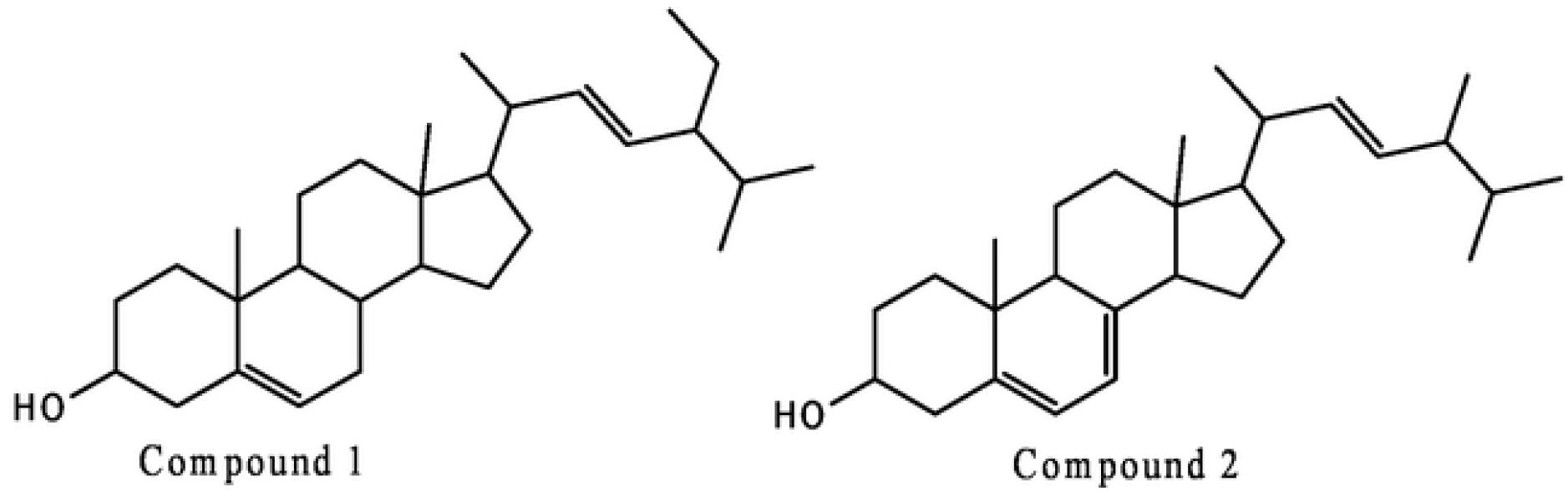
Structures of Compounds 1 and 2.

Nature: off-white crystalline powder

Yield: 73 mg

^1^H-NMR (500 MHz, CDCl_3_) and ^13^C-NMR (600 MHz, CDCl_3_). See (Table 5).

**Table 5:** Spectroscopic Data of Compound 1 (CDCl_3_)

| Position | Compound 1 |  | Stigmasterol (Literature) |  |
| --- | --- | --- | --- | --- |
| | $\delta_{\text{H}}$ (ppm) | $\delta_{\text{C}}$ (ppm) | $\delta_{\text{H}}$ (ppm) | $\delta_{\text{C}}$ (ppm) |
| 1 |  | 37.25 |  | 37.6 |
| 2 |  | 31.89 |  | 32.1 |
| 3 | 3.45, 1H, m, | 71.81 | 3.51, 1H, tdd | 72.1 |
| 4 |  | 42.46 |  | 42.4 |
| 5 |  | 140.75 |  | 141.1 |
| 6 | 5.34, 1H, s, | 121.71 | 5.31, 1H, t | 121.8 |
| 7 |  | 31.66 |  | 31.8 |
| 8 |  | 30.85 |  | 31.8 |
| 9 |  | 50.15 |  | 50.2 |
| 10 |  | 36.50 |  | 36.6 |
| 11 |  | 19.65 |  | 21.5 |
| 12 |  | 39.67 |  | 39.9 |
| 13 |  | 42.48 |  | 42.4 |
| 14 |  | 56.85 |  | 56.8 |
| 15 |  | 24.30 |  | 24.4 |
| 16 |  | 28.91 |  | 29.3 |
| 17 |  | 56.76 |  | 56.2 |
| 18 | 1.13, 3H, s | 12.40 | 1.03, 3H, s | 12.2 |
| 19 | 0.68, 3H, s | 18.91 | 0.71, 3H, s | 18.9 |
| 20 |  | 39.79 |  | 40.6 |
| 21 |  | 21.30 | 0.91, 3H, d, 6.2 Hz | 21.7 |
| 22 | 5.17, 1H, m, 8.2 Hz | 138.31 | 5.14, 1H, m | 138.7 |
| 23 | 5.00, 1H, m, 8.5 Hz | 129.26 | 4.98, 1H, m | 129.6 |
| 24 | 0.84, 1H, m, 6.7 Hz | 45.83 |  | 46.1 |
| 25 |  | 25.70 |  | 25.4 |
| 26 | 0.89, 3H, m | 12.25 | 0.83, 3H, t | 12.1 |
| 27 |  | 29.14 |  | 29.6 |
| 28 | 0.81, 3H, m | 19.90 | 0.82, 3H, d, 6.6 Hz | 20.2 |
| 29 | 0.80, 3H, m | 19.13 | 0.80, 3H, d, 6.6 Hz | 19.8 |

s,m,d,t,ddd means singlet, multiplets, doublets, triplets and triplets of doublets of doublets respectively

EIMS [M]^+^ m/z 421.2 (100) calculated for C_29_H_48_O, [M-CH_4_]^+^m/z 396.2 (43), [M-C_2_H_7_]^+^ m/z 381.2, 300.1 (41), 271.1 (54), 255.1 (78), 159.0 (42), 83 (40), 69 (27), 55.0 (41) 43 (20).

Compound **2** (Ergosterol) Figure 4

Nature: white powder

Yield: 45 mg

^1^H-NMR (500 MHz, CDCl_3_) and ^13^C-NMR (600 MHz, CDCl_3_) (Table 6).

**Table 6:** Spectroscopic Data of Compound 2 (CDCl_3_+CD_3_OD)

| Position | Compound 2 |  | Ergosterol (Literature) |  |
| --- | --- | --- | --- | --- |
| | $\delta_H$ (ppm) | $\delta_C$ (ppm) | $\delta_H$ (ppm) | $\delta_C$ (ppm) |
| 1 |  | 38.43 |  | 38.4 |
| 2 |  | 31.69 |  | 31.9 |
| 3 | 3.27, 1H, m, | 69.94 | 3.64, 1H, m | 70.5 |
| 4 |  | 39.50 |  | 42.8 |
| 5 |  | 138.7 |  | 139.9 |
| 6 | 5.27, 1H, s, | 121.91 | 5.58, 1H, dd, 5.5, 3.0 Hz | 119.6 |
| 7 |  | 100.85 | 5.38, 1H, dd, 5.4, 2.9 Hz | 116.3 |
| 8 |  | 140.05 |  | 141.4 |
| 9 |  | 45.61 |  | 46.2 |
| 10 |  | 37.00 |  | 37.1 |
| 11 |  | 19.47 |  | 21.1 |
| 12 |  | 40.01 |  | 39.1 |
| 13 |  | 39.50 |  | 42.9 |
| 14 |  | 55.81 |  | 54.6 |
| 15 |  | 22.27 |  | 22.9 |
| 16 |  | 28.89 |  | 28.3 |
| 17 |  | 56.51 |  | 55.7 |
| 18 |  | 11.65 | 0.95, 3H, s | 12.1 |
| 19 |  | 15.75 | 0.65, 3H, s | 16.3 |
| 20 |  | 40.41 |  | 40.3 |
| 21 | 1.01, 3H, s | 20.80 | 1.04, 3H, d, 6.6 Hz | 21.1 |
| 22 | 5.10, 1H, m, 8.1 Hz | 130.31 | 5.20, 1H, m | 135.6 |
| 23 | 4.97, 1H, m | 129.4 | 5.21, 1H, m | 131.9 |
| 24 | 0.84, 1H, m, 6.7 Hz | 43.18 |  | 42.9 |
| 25 |  | 33.69 |  | 33.1 |
| 26 | 0.88, 3H, m | 19.01 | 0.84, 3H, d, 6.7 Hz | 19.9 |
| 27 | 0.86, 3H, m | 18.68 | 0.82, 3H, d, 6.7 Hz | 19.7 |
| 28 | 0.93, 3H, m | 18.46 | 0.82, 3H, d, 6.6 Hz | 17.6 |

s,m,d,t,ddd,dd means singlet, multiplets, doublets, triplets and triplets of doublets of doublets, doublets of doublet respectively

EIMS [M]^+^ m/z 396.4 (100) calculated for C_28_H_44_O, [M-CH_2_]^+^ m/z 382 (23), 255 (21), 147 (19).

Spectroscopic identification of isolated compounds revealed that compounds **1** and **2** have a steroidal moiety. The mass spectrum of compound **1** showed a molecular ion peak [M]^+^ at m/z 421.2 which is also the base peak, followed by other fragment ions at m/z 396.2 [M-CH_4_]^+^which due to loss of methane and m/z 381.2 [M-C_2_H_7_]^+^. The molecular mass of 421.2 is consisted with molecular formula, C_29_H_48_O. The ^1^H-NMR spectrum of compound 1 revealed 3 downfield proton signals in the olefinic region at δ_H_ 5.34 (1H, s, 1.6Hz, H-6); 5.17 (1H, m, 8.2Hz, H-22) and 5.00 (1H, m, 8.5Hz, H-23). The 8.2Hz coupling constant indicating the *cis* nature of the olefinic double bond between C-22 and C-23, which was confirmed by the NOESY spectrum. The signal at δ_H_ 3.45 (1H, m, H-3) was confirmed by DEPT-90 spectrum to be a methine carbon (CH) bonded to a highly electronegative oxygen. The spectrum showed many alkyl signals in the upfield region between δ_H_ 0.5-2.5ppm which is due to stigmastane nucleus of Compound **1**. The ^13^C-NMR spectrum and DEPT-90 spectra of Compound **1** showed signals due to 4 non-equivalent sp^2^ hybridized olefinic carbons at δ_c_ 140.75 (C-5), 138.31 (CH-23), 129.26 (C-22) and 121.71 (CH-6). Also, a sp^3^ hybridized oxygenated methylene carbon at δ_c_ 71.81 (CH-3). The comparison of DEPT-135 with^13^C spectra further confirmed that carbon signal at δ_c_ 140.75 is a quaternary olefinic carbon. The stigmastane steroidal nucleus of Compound 1 is indicated by signals in the upfield region of the spectrum at δ_c_ 10-60 which is characteristic of the steroidal family. The COSY and NOESY spectra information agree with the literature data on Stigmasterol ^[19]^.

The mass spectrum of Compound **2** showed a molecular ion peak [M]^+^ at m/z 396.4 (100) which is also the base peak. Other fragment ions included [M-CH_2_]^+^m/z 382 (23) loss of CH_2_ unit, 255 (21), 147 (19). The molecular mass of 396.4 is consistent with molecular formula C_28_H_44_O. The molecular mass of compound 2 therefore showed a striking difference from compound **1**. The ^1^H-NMR of Compound **2** showed similar spectra information as obtained in **1** with olefinic protons at δ_H_ 5.27 (1H, s, 1.6Hz, H-6), 5.13 (1H, m, H-7), 5.00 (1H, m, H-22), 4.99 (1H, m, H-23). The signal at δ_H_ 3.27 (1H, m, H-3) is confirmed by the DEPT-90 experiment to be a methylene carbon (CH) bonded to a highly deshielding oxygen. The spectrum further showed many signals in the upfield region at δ_H_ 0.5-2.5 which is due to the steroidal hump of ergostane nucleus of Compound **2**. The ^13^C NMR spectrum and DEPT-90 spectra of Compound **2** showed signals due to 6 non-equivalent sp^2^ hybridized olefinic carbons at δ_c_ 140.05 (C-8), 138.71 (C-5), 130.33 (CH-22), 129.41 (CH-23), 121.91 (CH-6) and 100.89 (CH-7). Furthermore, a sp^3^ hybridized oxygenated methylene carbon at δ_c_ 69.97 (CH-3). The comparison of ^13^C and DEPT-135 spectra further indicated that carbon signal at δ_c_ 140.05 and 138.71 are quartenary olefinic carbon signals, which further differentiate compound **2** from **1**. The steroidal hump nucleus of **2** is indicated by many signals in the upfield region of the spectrum at δ_c_ 10-60 typically common to most steroids. The spectral information thus obtained on **2** is in agreement with literature data on Ergosterol ^[20].^

## 4. DISCUSSION

Brine shrimp lethality assay (BSLA) is a straightforward and inexpensive bioassay used for testing the cytotoxic effect of phytochemical present in the plant extracts [21]. The result on the lethality of methanolic extract and fractions of *S. biafrae* plant on brine shrimps showed that the plant extract and fractions exhibited a good toxicity with LC50 < 1000 µg mL. Based on the results of this study, it can be reported that the hexane and dichloromethane fractions of the plant demonstrated relatively lower toxicity compared to the other fractions or extract tested, as well as the standard drug (Doxorubicin). This suggests that these particular fractions may have safer properties and could potentially be explored further for their therapeutic applications with reduced risk of adverse effects. However, it is important to note that further research and evaluation are necessary to fully understand the safety profile and potential benefits of these fractions in the context of drug development and cancer management. According to Meyer *et al*. (1982), substances with LC50 value < 1000 µg mL are toxic while ones with values > 1000 µg mL are non-toxic. The observed toxic property of the n-hexane fraction against brine shrimp larvae provides additional support for its cytotoxic activity against the PC-3 cell line. The cytotoxicity exhibited by the n-hexane fraction against the brine shrimp larvae suggests its potential as a bioactive compound with cytotoxic effects. This finding supports the potential of the n-hexane fraction as a candidate for further investigation and development as an anticancer agent against the PC-3 cell line. The cytotoxic activity revealed that n-hexane fraction of the methanolic extract *S. biafrae* leaf exhibited strong inhibitory activity on PC-3 cell line, followed by the standard drug doxorubicin, dichloromethane, compound 1 and 2. Compound 1 was found to exhibit moderate % inhibition on the proliferation of HeLa and PC-3 cell lines respectively compared to the standard drug used. Compound 1 exhibited a better cytotoxic activity than compound 2. Thus, the higher cytotoxic activity exhibited by the *n*-hexane fraction could be attributed to the bioactive compounds that are present in the fraction and their synergistic interaction [23]. This strong inhibitory activity observed in hexane fraction against PC-3 cell line is in line with the previous studies on other *Senecio* species. According to Loizzo *et al*. [24], the n-hexane extract of *S. ambiguus* exhibited a strong inhibitory activity against prostate carcinoma cell line, a similar result to the n-hexane fraction of *S. biafrae* against prostate cancer cell line indicating that all *Senecio* species might have some activities in common.

The chemical profiling of hexane fraction from *S. biafrae* plant revealed the presence of seventeen (17) while DMF had six (6) saturated and unsaturated fatty acids, phytol and benzene dicarboxylate esters. The most abundant saturated fatty acid identified is n-hexadecanoic acid (palmitic acid) known for exerting anti-cancerous, anti-inflammatory, antioxidant and anti-androgenic pharmacological effects [25]. This was followed by unsaturated fatty acid cis, cis, cis-9,12,15-octadecatrienoic acid and phytol, an acyclic diterpene alcohol. Phytol is a known precursor of phytanic acid, vitamins E and K and has been found to possess strong cytotoxic, antioxidant and anti-inflammatory activities [26-27]. These compounds were characterized as white amorphous solids and exhibited a positive reaction with Salkowski reagents, indicating the presence of steroidal structures. Phytosterols are plant-derived compounds that have been widely studied for their various biological activities, including anticancer properties. The identification of these phytosterols in the plant extract suggests their potential contribution to the observed cytotoxic activity. The molecular formula of the isolated molecules found to be stigmasterol and ergosterol are C_29_H_48_O and C_28_H_44_O respectively which was confirmed by mass spectral and nuclear magnetic resonance of 1D and 2D structure. Phytosterols being a steroidal alcohol play important roles in membrane stability and in structural component in the cell membrane. Since pyhtosterol cannot be synthesized by human being or animal, they are derived through the exogeneous sources, especially the diet [28]. Phytosterols are very abundant in plants, they share many structural similarities with cholesterol which has been implicated in the pathophysiology of many oxidative stress-induced conditions. Phytosterols have been speculated to possess anti-cancer activities in many organs of the body such as ovary, lungs and stomach [29]. The high amount of phytosterol intake is from vegetarian diet; that are consistent with a reduction of breast cancer risk [30]. Increasing the concentration of phytosterols in the blood can indeed be achieved through the extraction of phytosterols from plant foods and their incorporation into various other food products or dietary supplements. Phytosterols are naturally occurring compounds found in plant cell membranes, and they have a similar structure to cholesterol [31, 32]. By consuming foods or supplements enriched with phytosterols, individuals can potentially reduce cholesterol absorption in the intestine, leading to lower LDL (bad) cholesterol levels in the blood [33]. Vegetables like *S. biafrae* can be a potential source of phytosterols, and their extraction and incorporation into different food products can provide a convenient way for individuals to increase their phytosterol intake. These enriched products may include fortified spreads, margarines, yogurts, beverages, and dietary supplements. Phytosterol-enriched products are commonly available in the market and are specifically marketed for their cholesterol-lowering benefits. Additionally, incorporating a variety of plant-based foods rich in phytosterols, along with a balanced diet and a healthy lifestyle, is essential for overall well-being.

## CONCLUSION

The results of the study showed that the methanolic extract fractions of *S. biafrae* leaf exhibited good toxicity when tested against brine shrimp larvae. The hexane fraction of *S. biafrae* leaf exhibited a significant cytotoxic activity on PC-3 cell than other fractions and isolated compounds. The cytotoxic activity observed by hexane fraction may be due to the presence of chemical constituents and their synergistic interaction. Also, the use of mass spectral, ^1^H NMR, ^13^C NMR including 2D-NMR analyses enabled the identification of two phytosterol compounds: stigmasterol and ergosterol in dichloromethane fraction of *S. biafrae*. Compound 1 (stigmasterol) showed a moderate activity on HeLa cell while other fractions and compound 2 (ergosterol) exhibited a weak activity.

## ACKNOWLEDGEMENTS

Akinola F.T. acknowledges the World Academy of Sciences for the Advancement of Science in Developing Countries (TWAS), Trieste, Italy for ICCBS-TWAS Fellowship at the H.E.J. Research Institute of Chemistry, ICCBS, University of Karachi, Karachi, Pakistan. The authors are also grateful to the MS and NMR Unit of H.E.J Research Institute of Chemistry, ICCBS, University of Karachi, Karachi, Pakistan for recording the GC-MS, NMR and MS spectrum of hexane fraction and isolated compounds respectively.

## REFERENCES

1. Cragg GM and Newman DJ. Biodiversity: A Continuing Source of Novel Drug Leads. Pure and Applied Chemistry 2005 77:7–24.

2. Firenzuoli F and Gori L. Herbal Medicine Today: Clinical and Research Issues. Evidenced-Based Complementary and Alternative Medicine 2007. 4:37–40

3. Heinrich M, Barnes J, Gibbons S and Williamson EM. Fundamentals of Pharmacognosy and Phototherapy. Churchill Livingstone: 2004. Elsevier Science Ltd., UK.

4. Nasreen S and Radha R. Assessment of Quality of Withania somnifera Dunal (Solanaceae) Pharmacognostical and Phyto-Physicochemical Profile. International Journal of Pharmacy Pharmaceutical Sciences 2011. 3: 152–155.

5. Panda SK, Thatoi HN and Dutta SK. Antibacterial Activity and Phytochemical Screening of Leaf and Bark Extracts of Vitex negundo from Similipal Biosphere Reserve Orissa. Journal of Medicinal Plant Research. 2009. 3:294–300.

6. Okigbo RN, Eme UE and Ogbogu S. Biodiversity and Conservation of Medical and Aromatic Plants in African. Biotechnology and Molecular Biology Reviews. 2008. 3:127–134.

7. UNESCO. Culture and Health, Orientation Texts – World Decade for Cultural Development 1988– 1997, Document CLT/DEC/PRO – 1996, Paris, France, 1996, pp 129.

8. Nascimento GGF, Locatelli J, Freitas PC and Silva GL. Antibacterial activity of plant extracts and phytochemicals on antibiotic-resistant bacteria. Brazilian Journal of Microbiology 2000.31:247–256.

9. Saikia B. Ethnomedical Plants from Gohpur of Sonitpur District, Assam. Indian Journal Traditional Knowledge 2006.5: 529–530.

10. Bello OA, Ayanda OI, Aworunse OS, Olukanmi BI, Soladoye MO, Esan EB et al. Solanecio biafrae: An underutilized nutraceutically-important African indigenous vegetable. Pharmacognosy Review. 2018.12:128–32.

11. Ogundele SB, Oriola AO, Oyedeji AO, Olorunmola FO, Agbedahunsi JM. Flavonoids from Stem Barks of Artocarpus altilis (Parkinson ex F. A. Zorn) Fosberg. Chemistry Africa, 2022. 1–15

12. Amir AM, Omid AK, Amir HS. Phytochemical Analysis and Cytotoxicity Evaluation of Kelussia odoratissima Mozaff. Journal of Acupuncture and Meridian Studies. 2017. 10(3):180e–186.

13. Olasupo AD, Olagoke OV and Aborisade AB. Qualitative Phytochemical Screening of Bologi (Senecio biafrae) and Bitter Leaf (Vernonia amygdalina) Leaves. Chemical Science International Journal. 2017. 20(3): 1–6.

14. Ajiboye BO, Ojo OA, Okesola MA, Akinyemi AJ, Talabi JY, Idowu OT et al. In vitro antioxidant activities and inhibitory effects of phenolic extract of Senecio biafrae (Oliv and Hiern) against key enzymes linked with type II diabetes mellitus and Alzheimer’s disease. Food Science and Nutrition. 2018. 1–8.

15. Adelakun S, Omotoso O, Aniah J and Oyewo O. Senecio biafrae defeated Tetracycline-Induced Testicular Toxicity in Adult Male Sprague Dawley Rats. Jornal Brasileiro de Reproducao Assistida, 2018a. 1-22(4):314–322.

16. Adelakun S.A, Ogunlade B, Omotoso O.D. and Oyewo, O.O. Role of Aqueous Crude Leaf Extract of Senecio biafrae Combined with Zinc on Testicular Function of Adult Male Sprague Dawley Rats. Journal of Family and Reproductive Health. 2018b. 12(1): 8–17.

17. Somaida A, Tariq I, Ambreen G, Abdelsalam AM, Ayoub AM. Potent Cytotoxicity of four Cameroonian Plant Extract on different Cancer Cell Line. Pharmaceuticals. 2020. 13:1–19.

19. Charturvedula VSP and Prakash I. Isolation of Stigmasterol and Sitosterol from the dichloromethane extract of Rubus suavissimus. International Current Pharmaceutical Journal. 2017. 1(9):239–242.

20. Alexandre TR, Lima ML, Galuppo MK, Mesquita JT, Nascimento MA, et al. Ergosterol isolated from the Basidiomycete Pleurotus salmoneostramineus affects Trypanosoma cruzi plasma membrane and mitochondria. Journal of Venomous Animals and Toxins including Tropical Diseases. 2017. 23(30):1–10.

21. Sandeep W, Mohan KK and Vijay RP. Brine Shrimp Lethality Assay of the Aqueous and Ethanolic Extracts of the Selected Species of Medicinal Plants. Proceedings 2019. 41: 1–12.

22. Meyer BN, Ferrigini RN, Putnam JE, Jacobsen LB, Nichols DE and McLaughlin JL. Brine shrimp: A convenient general bioassay for active plant constituents. Planta Med. 1982. 45:31 35.

23. Alcaraz MJ and Jimenez MJ. Flavonoids as anti-inflammatory agents. Fitoterapia 1988. 59: 25–38.

24. Loizzo MR, Tundis R, Statti GA and Menichini F. Jacaranone: A Cytotoxic Constituent from Senecio ambiguus subsp, ambiguus (Biv.) DC. against Renal Adenocarcinoma ACHN and Prostate Carcinoma LNCaP Cells. Arch Pharm Res. 2007:6:701–707

25. Choudhary D, Shekhawat JK and Kataria V. GC-MS Analysis of Bioactive Phytochemicals in Methanol Extract of Aerial Part and Callus of Dipterygium glaucum Decne. Pharmacogn J. 2019: 11(5):1055–1063

26. Islam MT, Ali ES, Uddin SJ, Shaw S, Islam MA, Ahmed MI, et al. Phytol: A review of biomedical activities. Food Chemical Toxicology. 2018. 121:82–94.

27. Islam MT, de Alencar MV, da Conceição Machado K, da Conceição Machado K, deCarvalho Melo-Cavalcante AA, de Sousa DP, et al. Phytol in a pharma-medico-stance. Chem Biol Interact. 2015. 240:60–73.

28. Kaur G, Gupta V, Singhai RGand Parveen B. Isolation and Characterization of Stigmasterol from Fritillaria roylei. Biology Medicine and Natural Product Chemistry. 2020. 9: 77–80

29. Bhagyashri V and Vinay S. Impact of different phytochemical classes and ayurvedic plants in battle against cancer. International Journal of Pharma Sciences and Research. 2016. 7(10):1–13

30. Cui X, Dai Q, Tseng M, Shu XO, Guo YT, Zheng W. Dietary pattern and breast cancer risk in the Shanghai breast cancer study. Cancer Epidemioogyl, Biomarkers and prevention. 2007.16:1443–1448.

31. Awad, A. B., Fink, C. S., and Williams, H. Phytosterols as anticancer dietary components: Evidence and mechanism of action. Journal of Nutrition, 2000. 130(9), 2127–2130.

32. Salman, H., Bergman, M., Bessler, H., & Punia, R. (2006). Effects of a plant sterol-enriched spread on biomarkers of endothelial dysfunction and low-grade inflammation in hypercholesterolaemic subjects. Scandinavian Journal of Clinical & Laboratory Investigation, 2006. 66(5), 415–424.

32. Panawan S, Watcharapong C, Sugunya M, Suwaporn L, Somsuda T. and Vijittra, L. Structures of Phytosterols and Triterpenoids with Potential Anti-Cancer Activity in Bran of Black Non-Glutinous Rice. Nutrients. 2015. 7: 1672–1687.

